# A KDM4A-PAF1-mediated epigenomic network is essential for acute myeloid leukemia cell self-renewal and survival

**DOI:** 10.1101/2020.10.30.361881

**Authors:** Matthew E Massett, Laura Monaghan, Shaun Patterson, Niamh Mannion, Roderick P Bunschoten, Alex Hoose, Sandra Marmiroli, Robert MJ Liskamp, Heather G Jørgensen, David Vetrie, Alison M Michie, Xu Huang

## Abstract

Epigenomic dysregulation is a common pathological feature in human hematological malignancies. H3K9me3 emerges as an important epigenomic marker in acute myeloid leukemia (AML). Its associated methyltransferases, such as SETDB1, suppress AML leukemogenesis, whilst H3K9me3 demethylases KDM4C is required for mixed lineage leukemia rearranged AML. However, the specific role and molecular mechanism of action of another member of KDM4 family, KDM4A has not previously been clearly defined. In this study, we delineated and functionally validated the epigenomic network regulated by KDM4A. We show that selective loss of KDM4A is sufficient to induce apoptosis in a broad spectrum of human AML cells. This detrimental phenotype results from a global accumulation of H3K9me3 and H3K27me3 at KDM4A targeted genomic loci thereby causing down-regulation of a *KDM4A*-*PAF1* controlled transcriptional program essential for leukemogenesis, distinct from that of KDM4C. From this regulatory network, we further extracted a *KDM4A-9* gene signature enriched with leukemia stem cell activity; the *KDM4A-9* score alone or in combination with the known *LSC1*7 score, effectively stratifies high-risk AML patients. Together, these results establish the essential and unique role of KDM4A for AML self-renewal and survival, supporting further investigation of KDM4A and its targets as a potential therapeutic vulnerability in AML.

## Introduction

Acute myeloid leukemia (AML) is an aggressive blood cancer affecting mostly adult and elderly patients. Growing evidence recognises that aberrant epigenetic-lead transcriptional regulation, such as hypermethylation at CpG islands (1) or H3K79me2 (2), and hypomethylation at H3K4me3 (3) and H3K9me3 (4) contributes to AML initiation/maintenance. Corroborating this notion, pharmacological epigenetic inhibitors, such as DNA methylase inhibitor, azacytidine (1, 5) have received regulatory approval for the treatment of myelodysplastic syndrome. Therefore, investigation of molecular mechanisms underpinning the epigenetic dysregulation in AML contributing to leukemogenesis is of importance, as it may uncover leukemic dependent transcriptional network(s), the core gene signature of which could be used as potential biomarkers for AML patient stratification and/or prognostic predication.

The H3K9 methyltransferases, such as SETDB1, have been shown to negatively regulate AML leukemogenesis (6). The previously observed association between global H3K9me3 hypomethylation in primary AML blasts and adverse outcome of patient prognosis (4) further suggests a potential role of H3K9me3 associated epigenetic modifying enzymes in AML. Cheung *et al*. further identified a H3K9me3 demethylase, KDM4C as a cofactor of PRMT1 involved transcriptional complex in mixed-lineage leukemia rearranged (MLLr) and MOZ-TIF2 AML (7). In addition, simultaneous knockout (KO) of all three members of Kdm4 family (*kdm4a/b/c)* in *B6.SJL* mice led to a strong attenuation of MLL-AF9 AML(8), indicative of roles for the Kdm4 family in murine myeloid leukemia. However, the roles and therapeutic tractabilities of other individual members of the KDM4 family in human AML are not well understood.

We previously performed a lentiviral knockdown (KD) screen targeting individual putative epigenetic regulators in 12 human AML cell lines representing different molecular subgroups of AML and found that depletion of one KDM4 family member, KDM4A, lead to significant suppression of leukemia cell proliferation (9). Substantial evidence establishes that KDM4A has different roles in normal tissue development compared to other members of the KDM4 family; it is amplified/overexpressed in various malignancies including AML and correlates with poor outcome in ovarian cancer (10, 11). Therefore, we tested the hypothesis that human KDM4A drives a distinct oncogenic mechanism compared to that known for KDM4C in human myeloid leukemia. Herein we demonstrate that selective loss of KDM4A is sufficient to induce AML cell death. This detrimental phenotype results from a global accumulation of epigenetic modifications, H3K9me3 and H3K27me3 at KDM4A targeted genomic loci thereby causing differential regulation of a *KDM4A*-mediated selective transcriptional program, including a minimum 9-gene signature enriched with leukemia stem cell (LSC) activity, which can effectively stratify high-risk patients. These findings support an essential and unique role of KDM4A for AML cell self-renewal and survival.

## Materials and Methods

### Reagents, plasmids and virus manufacture

Puromycin and IOX1^dev^ were purchased from Sigma-Aldrich (St. Louis, MO, USA). IOX1 was from Tocris (#4464). pLenti-HA-KDM4A wt and mut (H188A/E190A) were a gift from Dr Gary Spencer (CRUK Manchester Institute). Lentiviral constructs for KD experiments were purchased from Sigma-Aldrich and are listed in the Supplemental Table. Lentiviral and retroviral supernatants were prepared, and leukemic human and murine cells transduced with viral particles as previously described (9).

### Flow cytometry, apoptosis and cell cycle analyses and immunoblotting

Flow cytometry analyses were performed using an LSRII flow cytometer (BD Biosciences, Oxford, UK). Cell sorting were performed using an Aria III flow cytometer (BD Biosciences). Details of antibodies used in flow cytometry and immunoblotting/immunoprecipitation are in the Supplemental Table.

### Immunofluorescence staining

Immunofluorescence (IF) staining was carried out using Hendley-Essex 12 well glass microscope slides. 6 × 10^4^ cells per condition were incubated on a poly-L-lysine coated slide for 1 hr before being fixed in 4% formaldehyde in PBS. The cells were permeabilized in 0.5% Triton-X-100 PBS followed by 2 hrs of blocking in 5% BSA, 0.2% Triton-X-100 TBS. Primary antibody diluted 1:500 in blocking solution was applied and slides incubated overnight in a humidified chamber at 4°C. Primary antibody was removed using PBS 0.1% Tween 20 (PBST) before a 1 hr room temperature incubation in appropriate secondary antibody (1:500 dilution in blocking solution) The antibody was again removed by washing with PBST. Antifade mountant with DAPI reagent (Thermo Fisher #P36962) was applied to each sample and a coverslip applied. After drying the slides were sealed and images captured at 40x/100x magnification on the Zeiss Axioimager M1 Epifluorescence and Brightfield Microscope.

### Culture of cell lines and primary cells

Leukemia cell lines were from DMSZ (Braunschweig, Germany). All cell lines were grown in the recommended cell culture media at 37°C in 5% CO_2_. Murine and human primary AML and normal BM samples were cultured as described (12). Murine MLL-AF9 AML cells were leukemic BM cells extracted from a cohort of mice with MLL-AF9 AML established by Somervaille et al (13), and cultured in the conditional medium with mIL-3 (100ng/ml). All cytokines were purchased from PeproTech (London, UK).

### Colony forming cell assay

Colony forming cell (CFC) assay for murine cells was performed by plating 1000 cells on methylcellulose (MethoCult #M3434, Stem Cell Technologies). Colony Assay for human CD34^+^ HSPCs and AML patient cells were performed by plating 10000 cells and 3000 cells respectively on methylcellulose (MethoCult #H4434, Stem Cell Technologies). CFU-GM (Granulocyte/Macrophage), M(Macrophage), and E(Erythroid) colonies were assessed and counted 10 days after seeding.

### Murine transplantation experiments

Experiments using mice were approved by the local animal ethics review board and performed under a project licence issued by the United Kingdom Home Office, in keeping with the Home Office Animal Scientific Procedures Act, 1986. Non-obese diabetic. Cg-Prkdc scid Il2rgtm1Wjl/SzJ (NSG) mice were purchased from Jackson Laboratories (Bar Harbor, ME, USA) for transplantation as previously described (9). Briefly, indicated primary AML patient samples for xenograft transplantation were unfractioned primary blasts from our and Manchester biobank collections. Control or *KDM4A* KD human AML THP1 cells or primary AML patient blasts were FACS sorted 40 hours following lentiviral infection and immediately transplanted into sub-lethally irradiated (450cGy) NSG mice (10,000 THP1 cells or 10^6 primary AML cells) via tail vein injection.

### RNA isolation, quantitative PCR, RNA-seq and ChIP-seq

AML cells were transduced as previously described with two distinct lentiviruses for *KDM4A* KD (*KDM4A* KD#1 and *KDM4A* KD#2) and two for *PAF1* KD (Supplemental Table-2). A non-targeting control (NTC) lentivirus was used as a control. RNA was extracted from transduced cells 72 hr following puromycin selection using QIAshredder™ columns and the RNeasy Plus Microkit™ (Qiagen). RNA-seq libraries were produced using the TruSeq® stranded mRNA kit (Illumina) and sequenced using the Illumina NextSeq™ 500 platform. For ChIP-seq, DNA was purified using Diagenode's iPure kit v2 and libraries made using the TruSeq ChIP Library Preparation Kit according to the manufacturer’s instructions. FastQC was used to inspect and ensure the quality of sequencing data. Independent experiments of QPCR and ChIP-QPCR were carried out for RNA-seq and ChIP-seq validation. Details of RNA-seq and ChIP-seq data analysis and *KDM4A-9* gene signature construction methods are in the Supplemental Methods. RNA-seq and ChIP-seq data files are available in the Gene Expression Omnibus (GEO): GSE125376. For gene expression correlation and survival analyses, processed datasets were downloaded from public databases: (i) E-MTAB-3322 (7) (ii) GSE81299 (8) (iii) GSE6891 (14) (iv) GSE12417 (15) (v) GSE37642 (16) (vi) Vizome (17) (vii) COSMIC (18).

### Study Approval

Use of human tissue was in compliance with the ethical and legal framework of the United Kingdom’s Human Tissue Act (2004) and the Human Tissue (Scotland) Act (2006). Normal CD34^+^ mobilized HSPC surplus to requirements were from patients undergoing chemotherapy and autologous transplantation for lymphoma and myeloma. Their use was authorized by the Salford and Trafford Research Ethics Committee and, for samples collected since 2006, following the written informed consent of donors. Normal human BM was collected with informed consent from healthy adult male donors, with the ethical approval of the Yorkshire Independent Research Ethics Committee.

Primary human AML samples were from Manchester Cancer Research Centre’s Tissue Biobank (instituted with approval of the South Manchester Research Ethics Committee) and Paul O’Gorman Leukaemia Research Centre’s hematological cell research biobank (with approval of the West of Scotland Research Ethics Committee 4). Their use was authorized following project review by the Research Tissue Biobank’s scientific sub-committee, and with the informed consent of donors.

### Statistical analysis

Survival probabilities were estimated using the Kaplan-Meier method (survival 2.43-3 and survminer 0.4.3 R packages) and compared with the log-rank test. To dichotomize patients as either having a high or low signature score a median cut-off was utilized. For the differential expression of *KDM4A-9* genes between KDM4A^high^ and KDM4A^low^ patients, Welch t-test was used. Normally distributed groups were compared using two-tailed student *t*-test, unless stated otherwise. Correlation analysis was performed using Pearson’s correlation.

Statistics were calculated using R-3.6.1. Statistical significance of differential gene expression was assessed by Welch’s t-test unless otherwise stated. For RNA-seq, differential expression analysis was performed using the DESeq2 1.26.0 R package.

## Results

### KDM4A is required for the survival of human and murine AML cells

Data from COSMIC (18) show that up to 3.62% AML patient samples were found to exhibit elevated KDM4A expression (Fig. S1A), and its expression is significantly different from those of other KDM4 family members (Fig. S1B), indicative of a distinct role. Although there is no significant association of *KDM4A* expression with any other existing risk factors of AML (data not shown), such as specific oncogenic mutations or cytogenetic features, its expression is found highly enriched in AML-LSC^+^ populations (Fig. S1C), suggesting that KDM4A may be important for maintenance of the LSC pool, the presence and size of which is negatively correlated with AML patient survival.

To define a distinctive functional requirement of KDM4A in human AML cells, we performed lentivirus shRNA KD of *KDM4A*, *KDM4B* and *KDM4C* in human AML THP1 cells. THP1 cells driven by MLL fusion gene, have high levels of expression of KDM4A and have previously been used in our targeted depletion screen of chromatin regulatory genes (9). Furthermore, two previous studies of MLLr-driven leukemia had examined the function of other KDM4 family members (7, 8) allowing us to assess and compare the role of KDM4A in this AML subtype. *KDM4A* KD THP1 cells exhibited the greatest decrease in cell proliferation compared to NTC cells, using KD of *MLL* and *MEN1*, as positive controls (Figs. 1A-1C). Consistent with previously published work, lentiviral KD of *KDM4C* also had an inhibitory effect on AML cell proliferation (7), while *KDM4B* KD had no effect (Figs. 1A). Substantial loss of colony forming cell (CFC) potential in methylcellulose assays was correlated with the expression of *KDM4A* in a dose-dependent manner when five distinct *KDM4A* KD shRNA targeting constructs were compared (Fig. 1D). *KDM4A* KD induced apoptosis (Figs. 1E & S1D) rather than cell cycle arrest (Fig. S1E), with a small but consistent increase of myeloid cell surface markers CD13 and CD86 (19) expression relative to NTC (Fig. S1F), suggesting that loss of KDM4A may promote myeloid differentiation in THP1 cells. The requirement of KDM4A for the survival of AML cells was further confirmed in primary MLLr-AML patient blasts (Figs. 1F & 1G) and murine MLL-AF9 AML cells (Fig. 1H). Importantly, we determined the impact of *KDM4A* KD on AML initiation *in vivo* by transplanting *KDM4A* KD THP1 cells (Fig. 1I) or primary MLLr-AML cells (Figs. 1J-1K & S1G-S1I) into recipient NSG mice. Control (NTC) cells induced short latency disease within 40 days with splenomegaly (Fig. 1K). Loss of KDM4A significantly prolonged overall survival of mice with only one mouse succumbing to leukemia over the follow-up period by either *KDM4A*#1 KD or *KDM4A*#2 KD (Figs. 1J-1K & S1G-S1I). Taken together, these data demonstrate a specific and essential role for KDM4A in AML cell survival.

**Figure 1.**
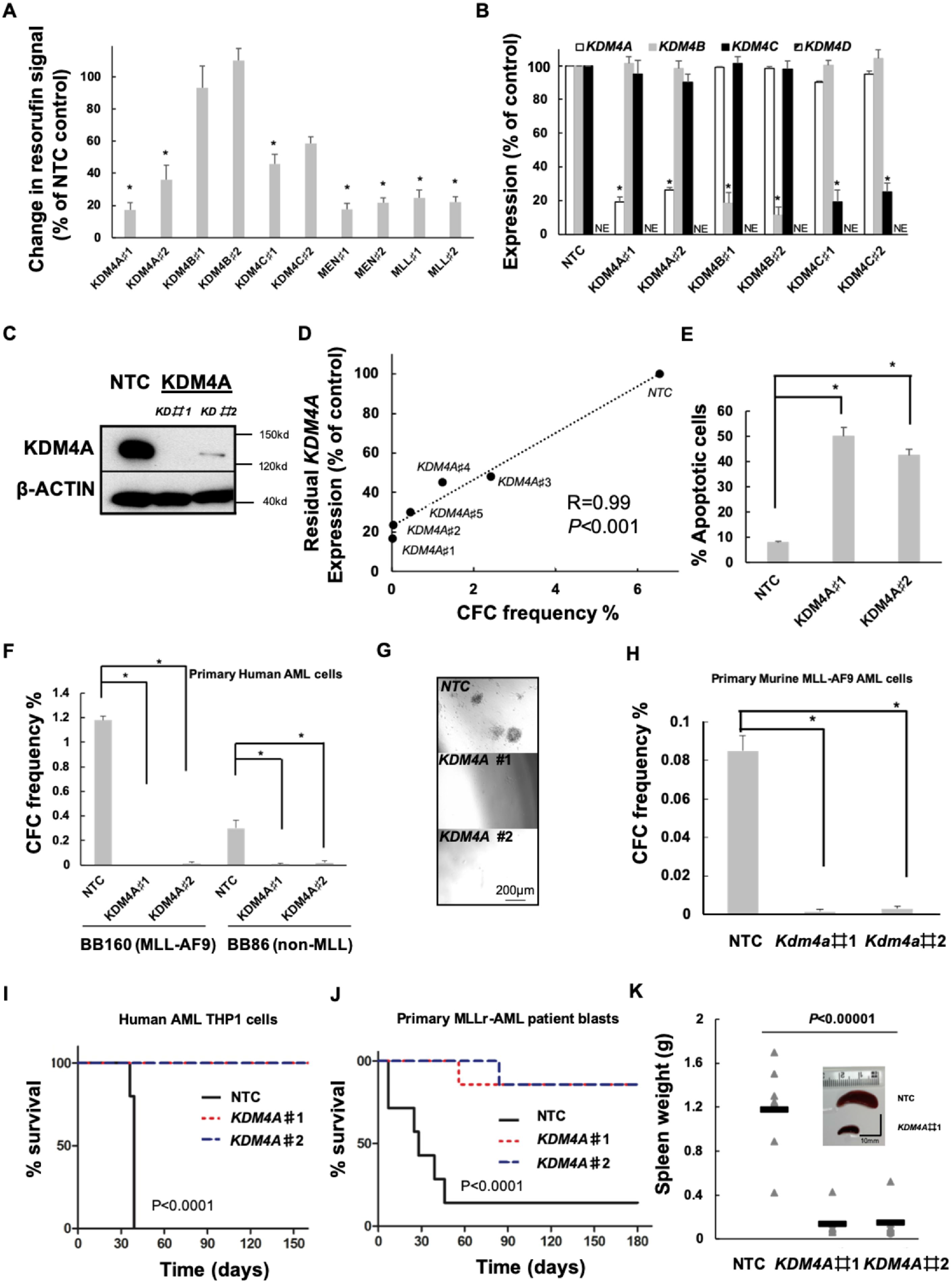
KDM4A is required for the functional potential of human and murine AML cells. (A-E) Human THP1 AML cells were transduced with lentiviruses targeting *KDM4A* or other KDM4 family members for KD (#1 & #2 represent individual distinct lentiviruses targeting genes for KD as indicated), or a non-targeting control (NTC). All bar charts show mean ± s.e.m.. (A) Resorufin signal after 4 days of individual KDM4 family member KD relative to NTC control cells (n=3); \**p*<0.01 for comparison of each KD versus *NTC*. (B) Expression of *KDM4A/B/C/D* in indicated KD cells relative to NTC control cells (n=3); \**p*<0.001. (C) Representative immunoblot showing *KDM4A* KD in THP1 cells (n = 3). (D) Scatter plot shows correlation of *KDM4A* KD with inhibition of frequency of colony forming cells (CFC) enumerated following 10 days in semisolid culture (n=3), as determined by QPCR; \**p*<0.001. (E) Percentage of apoptotic cells determined by Annexin V^+^/ 7AAD^+/-^ staining on day 4 of liquid culture after puromycin selection (n = 3); \**p*< 0.001. (F-G) The indicated primary unfractioned patient blasts were transduced with lentiviruses targeting *KDM4A* for KD, or an NTC. Primary AML cells used include BB160, containing t(9;11) (MLL-AF9) chromosomal translocation and BB86 (normal cytogenetics, non-MLL) (BB number is the Manchester Cancer Research Centre Biobank sample identifier). All bar charts show mean ± s.e.m. (F) CFC frequencies of primary human AML blasts (n=3) following lentivirus infection, puromycin selection and initiation of *KDM4A* KD; \**p*<0.0001. (G) Representative images from (F). (H) CFC frequencies of primary murine MLL-AF9 AML cells following *KDM4A* depletion (n=3); \**p*<0.0001. (I) Survival curves of NSG mice transplanted with 10,000 *KDM4A* KD or NTC THP1 cells (n=5 per cohort); *p* by log-rank test. (J) Survival curves of NSG mice transplanted with 10^6^ *KDM4A* KD or NTC primary AML cells (BB160, n=7 per cohort); *p* by log-rank test. (K) Spleen weights of mice from (J) with a representative image of the spleen. *p* by one-way ANOVA, F=34.13045.

### Targeting KDM4A’s demethylase activity inhibits AML cell proliferation

Next, we wanted to determine whether the catalytic demethylase activity of KDM4A is required for AML cells, we performed functional rescue experiments using murine MLL-AF9 cells. Forced-expression of wild-type human KDM4A rescued the clonogenic activity of AML cells transduced with *kdm4a* KD virus (Figs. 2A & 2B). However, this rescue phenotype was not observed when an enzymatically inactive mutant of human KDM4A (KDM4A^H188A/E190A^)(20, 21) (Figs. 2A & 2B) was expressed in murine MLL-AF9 cells. We further directly assessed the ability of KDM4A demethylase activity to modulate levels of H3K9me3 and H3K36me3 by examining the global changes of these established KDM4A substrates as readouts in *KDM4A* KD THP1 cells. As expected, there was a marked increase in the H3K9me3 level shown by immunoblotting (Figs. 2C & 2D) and an accumulation of H3K9me3 signal shown by immunofluorescent (IF) staining (Fig. S2A). No significant changes in H3K36me3 were observed by either approach (Figs. 2C-2D; S2A), suggesting H3K9me3 as the primary target of KDM4A in THP1 cells. This finding is in line with our ChIP-seq analysis of H3K36me3 changes in *KDM4A* KD THP1 cells (data not shown) and murine *kdm4a/b/c* triple KO cells (8). Additionally, there was a marked elevation of H3K27me3 levels globally in conjunction with the increase of H3K9me3 in *KDM4A* KD THP1 cells (Figs. 2C-2D; S2A). The global up-regulation of H3K9me3 and H3K27me3 was demonstrated in two further *KDM4A* KD human MLLr-AML cell lines, MV4:11 and MOLM13 (Fig. S2B).

**Figure 2.**
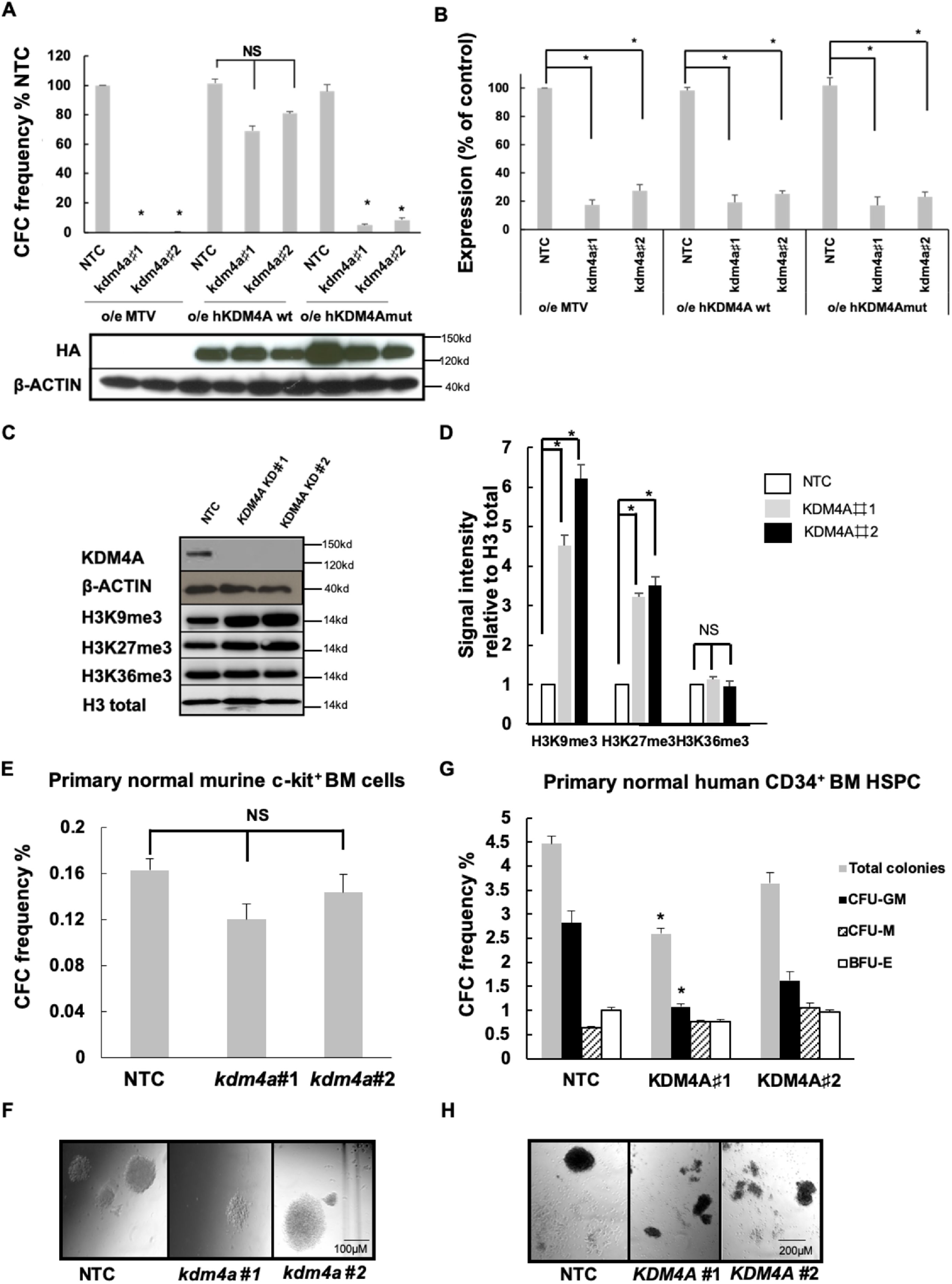
Targeting KDM4A’s demethylase activity inhibits AML cell proliferation. (A) CFC frequencies for control and *kdm4a* KD cells from the indicated murine MLL-AF9 cells overexpressing empty vector (MTV) or wild type human HA tagged-KDM4A or an enzymatically inactive mutant of human HA tagged-KDM4A (KDM4Amut H188A/E190A) (n=3); \**p*<0.001, ^NS^*p*>0.05. Representative immunoblot below bar plot shows the over-expression of wild type (wt) and mutant (mut) human HA-tagged KDM4A in correlated MLL-AF9 cells labeled, detected by HA antibody. (B) Bar chart showing mean ± s.e.m. expression of *kdm4a* by QPCR in *kdm4a* KD cells from (A) relative to NTC in murine MLL-AF9 leukemic cells (n=3); \**p*<0.01. (C) Representative immunoblots with indicated antibodies showing expression of indicated proteins in THP1 cells 72 hours following initiation of *KDM4A* KD (n=3). (D) Immunoblot quantification of signal intensity relative to H3 total from (C). (E-H) The indicated primary human and murine AML cells were transduced with lentiviruses targeting *KDM4A* or *kdm4a* for KD, or an NTC. All bar charts show mean ± s.e.m. CFC frequencies of (E) primary normal murine c-kit^+^ BM cells for NTC and *kdm4a* KD (n=3) or (G) primary normal human CD34^+^ HSPC cells for NTC and *KDM4A* KD cells (n=3); \**p*< 0.01. (F) and (H) are representative images from (E) and (G), Scale bar represents 100 μm and 200μm, respectively.

Inhibition of KDM4A’s demethylase activity resulted significantly impaired AML cell proliferation, making it an attractive therapeutic target. In furtherment of this idea, we investigated whether KDM4A is dispensable for normal hematopoiesis. The Kdm4 family are required for murine hematopoiesis in C57Bl/6 mice with *kdm4a/b/c* triple KO bone marrow (BM) cells unable to maintain functional hematopoiesis (8, 22). Normal hematopoiesis however can tolerate loss of a single Kdm4 family member as indicated in murine KO experiments (22). Consistently, our data confirmed no significant loss of colonies in *kdm4a* KD normal murine BM c-kit^+^ cells in methylcellulose assays (Figs. 2E & 2F; S2C). Methylcellulose assays using human CD34^+^ HSPCs from normal healthy donors demonstrate that reduced levels of *KDM4A* are generally tolerated (Figs. 2G & 2H; S2D) with less total number of colonies due to a noticeable reduction of CFU-GM in *KDM4A* KD #1 cells.

While a KDM4A specific inhibitor has not yet been reported, there are a number of KDM4 inhibitors in development including IOX1(23) and IOX derivatives (IOX1^dev^, *n*-Octyl ester-8-hydroxy-5-quinolinecarboxylic acid) (24, 25) as pan-KDM4 inhibitors. Similar to what was observed in other cancer cells (23, 24) (26), IOX1 and IOX1^dev^ displayed significant inhibition of cell proliferation in THP1 cells and primary AML patient blasts, inducing differentiation and apoptosis (Figs. S3A-S3G) with minimum effect on normal human CD34^+^ BM HSPCs (Fig. S3E). Importantly, these phenotypes were accompanied by an increased level of H3K9me3 and H3K27me3 (Fig. S3H), further supporting the anti-leukemic effect was a consequence of KDM4A demethylase inhibition. These results suggest KDM4A activity can be readily manipulated to compromise AML cell survival, supporting further investigation of KDM4A as a potential therapeutic vulnerability in AML.

### PAF1 identified as a KDM4A co-regulator is required for human AML cell survival

We next sought to define the molecular mechanism of KDM4A inhibition in killing AML cells. Profound epigenomic changes observed in *KDM4A* KD AML cells, indicate a significant transcriptional consequence following KDM4A depletion, causally related to its functional requirement in AML cell survival. To determine a KDM4A-maintained transcriptional network essential for AML cells, we compared the global transcriptome of *KDM4A* KD THP1 cells compared with NTC control cells by RNA-seq. We identified 3375 differentially expressed (DE) genes that are significantly deregulated following depletion of *KDM4A* compared with NTC (Log_2_ fold change (FC) ≥0.5 or ≤-0.5; adj. *p*≤0.05; Fig. 3A; supplemental file). Of these DE genes, 67% (2247 out of 3375) were direct targets of KDM4A; ChIP-seq revealed that KDM4A bound at their TSS (supplemental file). Given the fact that enriched H3K9me3(27) and H3K27me3(28) marks are often associated with heterochromatin formation leading to transcriptional repression, we would expect a significant down-regulation of KDM4A direct targets following its depletion. Indeed, 61% (1315 out of 2274) of putative KDM4A target genes were down-regulated, while 849 genes (39%) were up-regulated upon *KDM4A* KD in THP1 cells (Fig. 3A).

**Figure 3.**
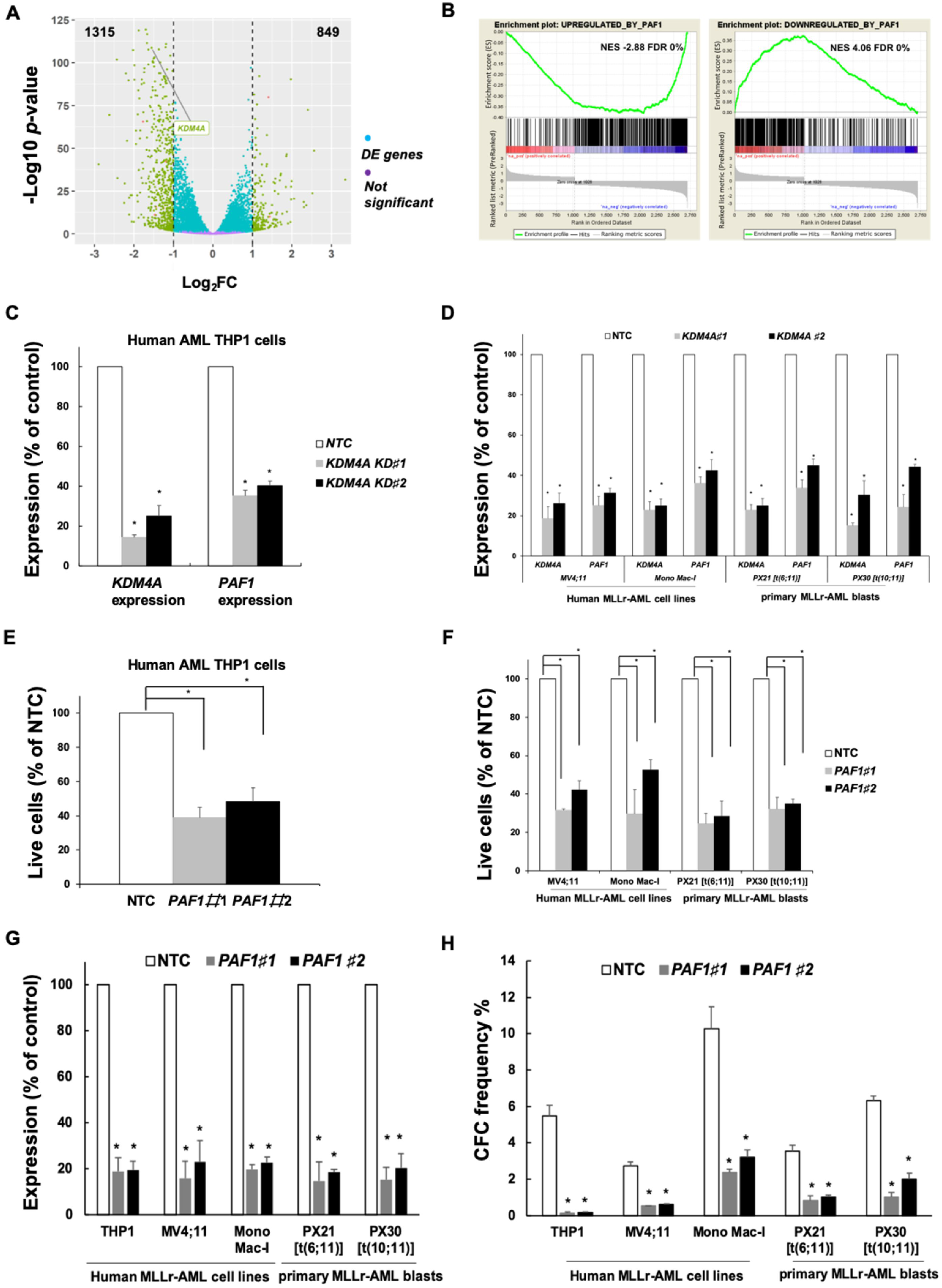
*PAF1* identified as a cofactor of KDM4A in MLLr-AML. (A) Volcano plot showing global changes in gene expression following loss of *KDM4A* compared to NTC control THP1 cells as identified by RNAseq. The absolute number of upregulated or downregulated genes which are bound by KDM4A are indicated at the top right and left side of the plot, respectively (KDM4A bound: FDR ≤ 0.01, gene expression: log_2_ FC ≥0.5 or ≤-0.5; adjusted (adj. *p*) *p*≤0.05). (B) GSEA show overlapping transcriptional consequences following loss of *KDM4A* or *PAF1* in THP1 cells. Specifically, genes repressed (left panel) or activated (right panel) are upregulated and downregulated respectively, following loss of KDM4A (29). (C-H) THP1 cells and other indicated human AML cells were transduced with lentiviruses targeting *KDM4A* or *PAF1* for KD, or an NTC. All bar charts show mean ± s.e.m. (C) Bar chart showing relative expression of *KDM4A and PAF1* by QPCR in comparison with NTC control cells following *KDM4A* KD using two different shRNA constructs *#*1 & *#*2 in THP1 cells (n=3) (D) and the other indicated human AML cells and primary AML cells include PX21, containing t(6;11)(MLL-AF6) chromosomal translocation and PX30 t(10;11)(MLL-AF10) (PX number is the Paul O’Gorman Leukaemia Research Centre Biobank sample identifier) (n=3) (F); \**p*< 0.01. (E-F) Percentage of live cell counts in comparison with NTC control (E) in THP1 cells 4 days following lentiviral infection (n=3) and (F) in the indicated human MLLr-AML cell lines and AML primary cells following *KDM4A* KD in comparison with NTC control cells (n=3) (I) \**p*<0.01. (G) Bar chart showing relative expression of *PAF1* by QPCR in comparison with NTC control cells following *PAF1* KD using two different shRNA constructs *#*1 & *#*2 in (E & F). (H) Bar chart showing the loss of CFC frequencies of indicated human AML cell lines and primary patient samples (n=3) following lentivirus infection, puromycin selection and initiation of *PAF1* KD; \**p*<0.001.

To provide insights into the survival pathways regulated by KDM4A, we performed gene-set enrichment analysis (GSEA) on our RNA-seq dataset and revealed a significant enrichment of genes regulated by polymerase associated factor 1 complex (PAF1c) (29–31)(Fig. 3B). This is consistent to the down-regulation of *PAF1* following *KDM4A* KD at transcript (Figs. 3C & 3D) and protein (Fig. S4A) level in human MLLr-AML cell lines, and primary MLLr-AML patient blasts. PAF1 is a core subunit of PAF1c that is essential for the proliferation of various subtypes of AML including those driven by MLLr fusions (32, 33) and has been implicated in a variety of solid tumours as well (34). Indeed, *PAF1* KD phenocopied *KDM4A* KD in MLLr-AML cells, inducing significant apoptosis (Figs. 3E-3G; S4B) and loss of CFU potential (Fig. 3H). Taken together, these data suggest loss of KDM4A impairs PAF1 function to maintain leukemic cell survival, supporting PAF1 as an important cofactor of KDM4A in human AML.

### KDM4A-PAF1 maintains appropriate expression of the MLLr-fusion oncogenic program in MLLr-AML

Corroborating the findings above, our ChIP-seq data show a substantial overlap amongst PAF1c (29), MLL-AF9 (35) and KDM4A binding sites in THP1 cells (Figs. 4A-4C; supplemental file). Specifically, KDM4A bound the PAF1 promoter region (supplemental file), suggesting a direct transcriptional regulatory mechanism. Further ChIP-seq analysis show that there is no significant enrichment of either histone methylation mark at non-KDM4A binding genomic loci (Figs. 4D & 4E), indicative of a human KDM4A-specific epigenomic profile. In marked contrast, using H3K9me3/H3K27me3 antibodies show a global gain of both H3K9me3 and H3K27me3 upon *KDM4A* KD in THP1 cells at KDM4A binding sites identified in control cells (Figs. 4F & 4G; 5A), including the genomic loci of PAF1 and its targets (Fig. 5B). These results were further validated by ChIP-QPCR in human MLLr-AML cell lines (Fig. 5C) and primary patient blasts (Fig. 5D).

**Figure 4.**
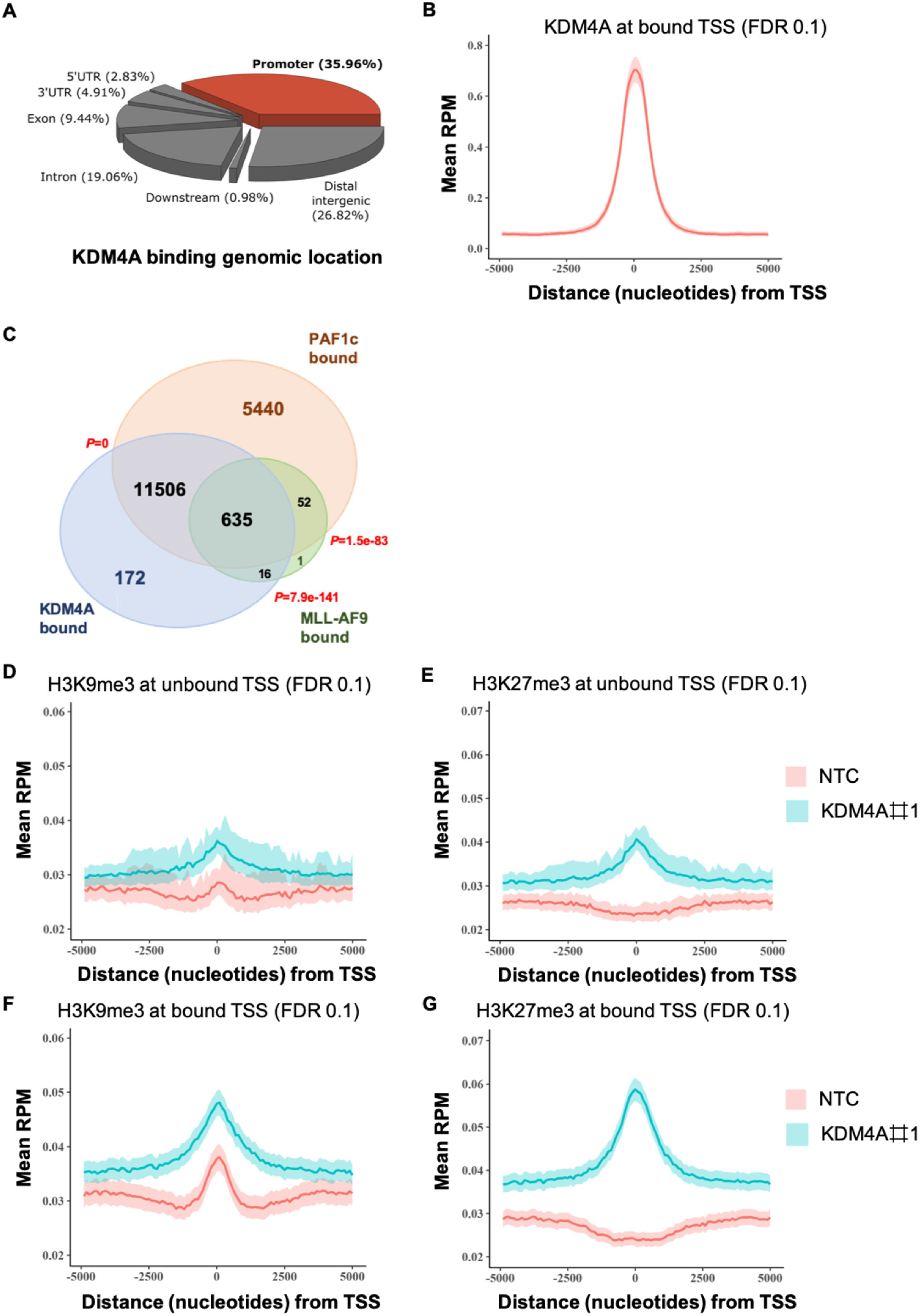
KDM4A-PAF1 co-regulates essential MLLr-fusion oncogenic transcriptional program. (A) Feature distribution of KDM4A ChIP-seq peaks in the THP1 cell genome. (B) Metagene plots showing a distinct peak in KDM4A normalised ChIP-seq signal in reads per million mapped reads (RPM) at transcription starting sites (TSS) in WT THP1 cells. (C) Venn diagram showing the overlap between binding sites of KDM4A, PAF1c (29) and MLL-AF9 in THP1 cells as determined by ChIP-seq; *p* by hypergeometric test. (D-G) Metagene plots showing an enrichment of H3K9me3 (F) and H3K27me3 (G) at KDM4A bound TSS compared to unbound TSS (D & E) following *KDM4A* KD by ChIP-seq.

**Figure 5.**
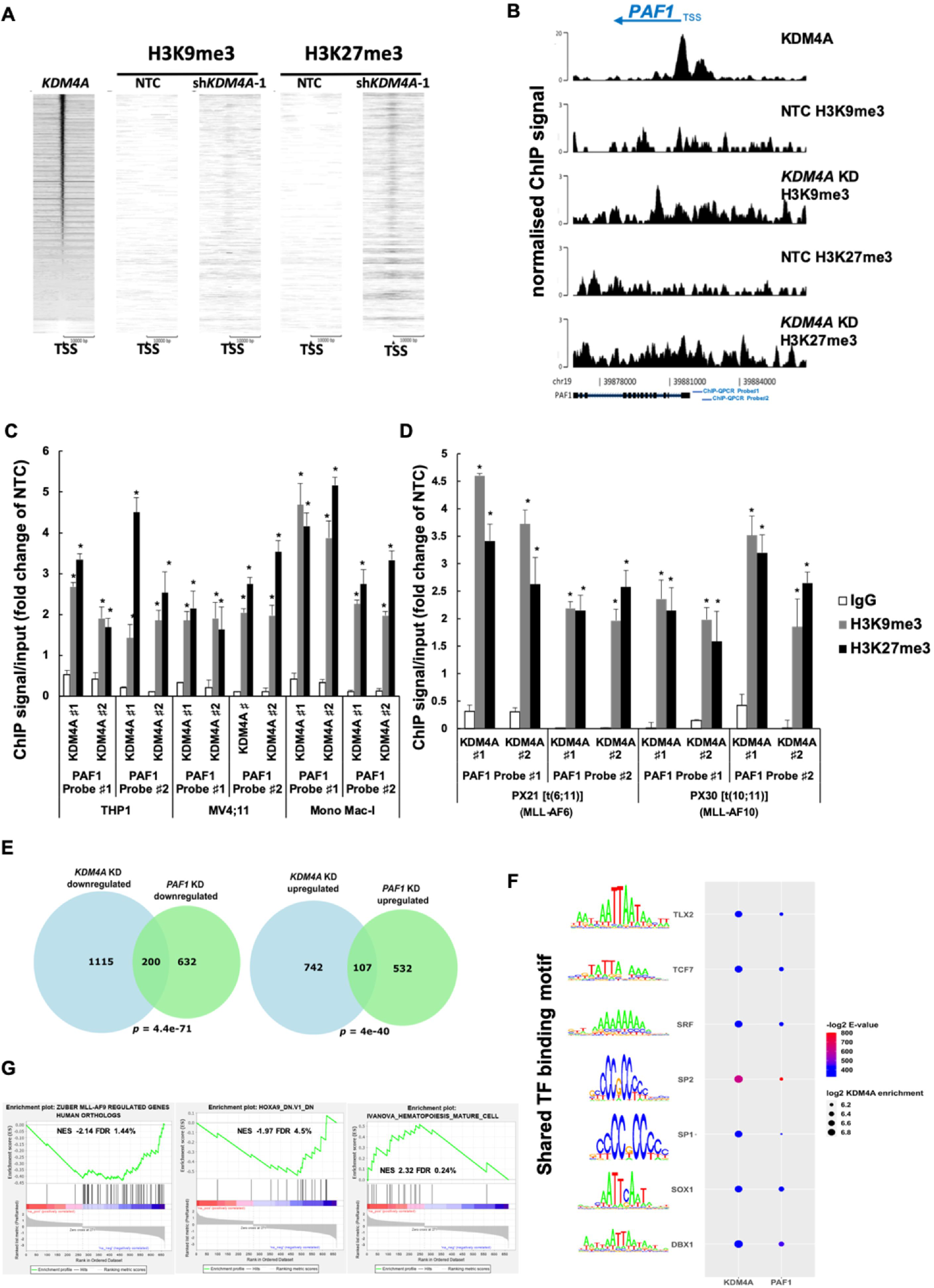
KDM4A-PAF1 maintains appropriate expression of the MLLr-fusion oncogenic program in MLLr-AML. Heatmap showing normalised ChIP-seq signal of H3K9me3 and H3K27me3 at TSS across all genes in *KDM4A* KD and NTC THP1 cells ordered by KDM4A enrichment. (B) Genomic snapshot demonstrates KDM4A occupancy at the PAF1 promoter region and enrichment of H3K9me3 and H3K27me3 signal throughout the PAF1 gene body and promoter upon *KDM4A* KD in comparison with NTC control in THP1 cells. Blue bars show the two individual probes used for ChIP-QPCR in (C & D). (C-D) H3K9me3 and H3K27me3 ChIP signal/input (fold change of NTC) in the indicated human MLLr-AML cell lines (C) and indicated primary AML samples including PX21 (MLL-AF6) and PX30 (MLL-FA10) (D) as determined by ChIP-QPCR following depletion of *KDM4A*; \**p*<0.001. (E) Venn diagrams showing the overlap between directly bound downregulated and upregulated targets of KDM4A and the PAF1c (24) in THP1 cells following knockdown of *KDM4A* and *PAF1* as determined by ChIPseq; *p* by hypergeometric test. (F) Motif significance and KDM4A log_2_ enrichment at KDM4A or PAF1 regulated promoters (FDR≤0.01, DE ≤ −0.5 or DE ≥ 0.5). Color represents motif significance within KDM4A and PAF1 regulated promoters. Size denotes the average log_2_ enrichment of KDM4A within each group of promoters that possess the respective transcription factor (TF) binding motif. Top five motifs detected in KDM4A or PAF1 regulated promoters sorted by statistical significance (E-value). (G) GSEA results showing significant overlap of *KDM4A* KD transcriptional consequences with down-regulation of MLL-AF9 and HOXA9 targets and up-regulation of a mature hematopoiesis program in THP1 cells, \**q*< 5%.

Furthermore, genes with significant expression changes following KDM4A silencing were also significantly enriched in direct PAF1 target genes (29)(Figs. 5E; supplemental file), suggesting a transcriptional network co-regulated by both KDM4A and PAF1. This notion is supported by the motif analysis of the promoter regions of KDM4A or PAF1 directly regulated genes (supplemental file), showing that KDM4A bound promoters share almost identical enrichment of transcription factor (TF) binding motifs as the ones bound by PAF1, including notably homeobox (HOX) transcription factors, such as TLX2 and DBX (Fig. 5F; supplemental file; 96%, E-value ≤0.05). Further GSEA analysis on the overlapped DE genes between *KDM4A* KD and *PAF1* KD revealed a significant downregulation of MLLr fusion target genes (36) as well as HOX family target genes (37) including notably the pro-survival gene, *BCL2*, and a marked upregulation of a mature hematopoiesis program (38) consistent to the differentiation phenotype observed upon *KDM4A* KD (Fig. 5G), such as *JUN* and *GATA2* and pro-apoptotic gene, *BCL2L11* (*BIM*). Although expression of *HOXA9* itself was not affected by either KD, our data suggest KDM4A and PAF1 co-regulate their downstream targets in a parallel manner. Collectively, we demonstrate that KDM4A collaborating with PAF1 plays a critical role in controlling essential gene expression network required for MLLr-AML cell survival via epigenomic regulation of H3K9me3 and H3K27me3.

### A core 9-gene signature downstream of *KDM4A* strongly associated with LSC activity and clinical outcome

Supporting the critical and collaborative role of KDM4A-PAF1 in AML, *KDM4A* expression is highly associated with *PAF1* expression in large AML patient datasets representing different subtypes (Figs. 6A & 6B); *KDM4A-PAF1* expression together can identify patients with inferior overall survival (Fig. 6C). These evidences suggest that KDM4A is required to sustain the survival and functional potential of AML cells across a broad spectrum of subtypes, rather than being confined solely to the MLLr molecular subtype. Consistently, *KDM4A* KD induced significant reduction of cell proliferation were found in additional human AML cell lines representative of different molecular subtypes (Fig. S5A), coupled with an increase in apoptosis and loss of CFC potential (Figs. S5B & S5C).

**Figure 6.**
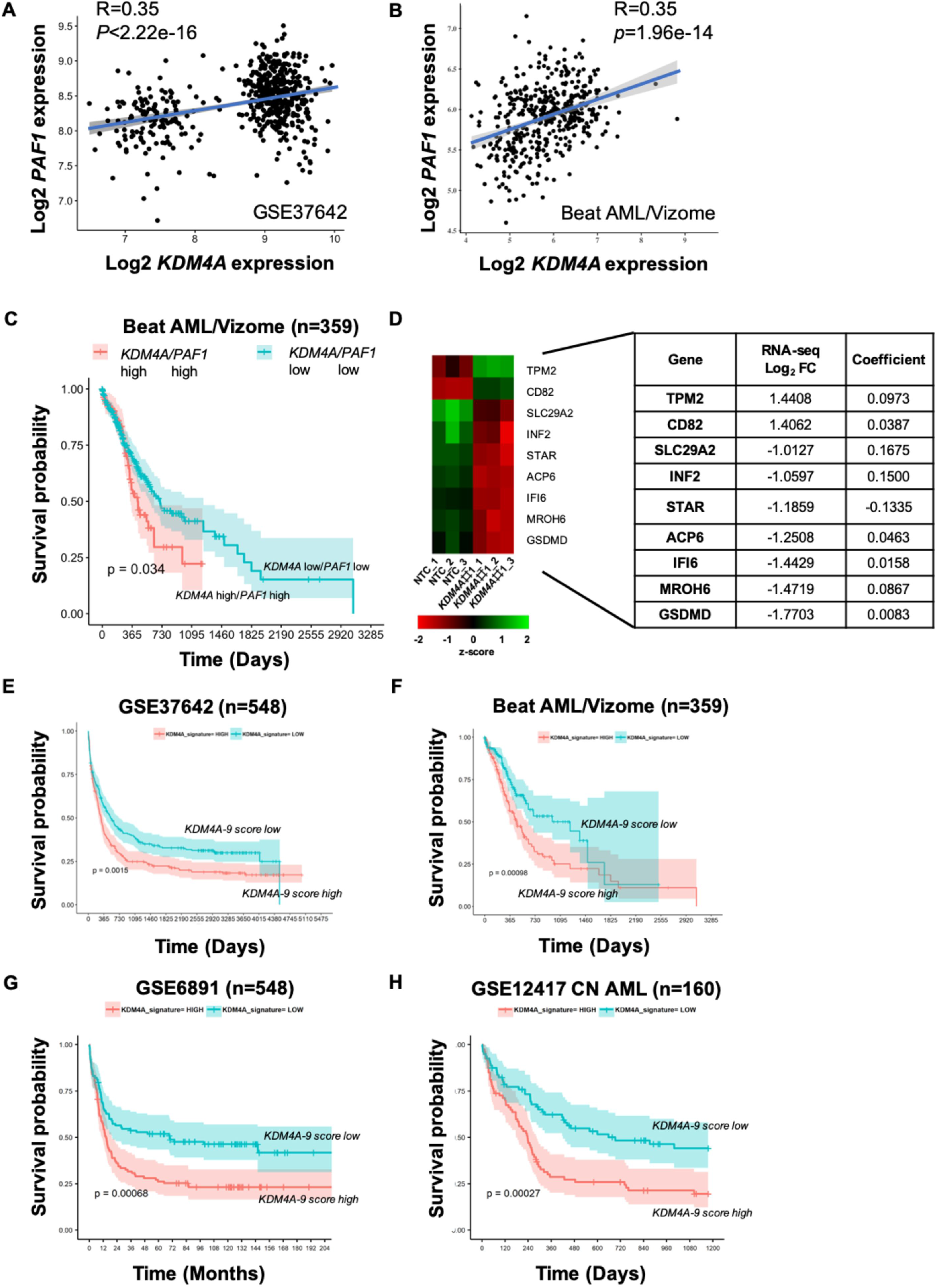
A core 9-gene signature downstream of *KDM4A* strongly associated with clinical outcome. (A-B) Scatterplot showing the correlation between expression of *KDM4A* versus *PAF1* in primary AML patient samples (GSE37642) (A) and (Beat AML/Vizome), R by Pearson correlation, *p*<0.05. (B). (C) Kaplan-Meier survival analysis conducted in Beat AML dataset. Patients with both *KDM4A*^high^ and *PAF1*^high^ expression have inferior overall survival. Patients dichotomized into high and low groups for *KDM4A* or *PAF1* based on whether expression was above the median for each gene; *p* by log-rank test. (D) Heatmap showing gene expression of the *KDM4A-9* gene signature genes with the table of their respective regression coefficients and log_2_ FC as determined by RNA-seq in *KDM4A* KD THP1 cells. (E-H) Kaplan-Meier survival analysis conducted in the large AML datasets (GSE37642) (E), (Beat AML/Vizome) (F), (GSE6891) (G) and (GSE12417) (H) showing that the *KDM4A-9* score can predict survival across AML patients of varying subtypes. Patients were dichotomized into high and low groups based on whether they possessed a score above or below the median signature score; *p* by log-rank test.

These lead to our hypothesise that there is a core gene signature downstream of *KDM4A-PAF1* regulatory axis, which could be extracted, specifically associated with patient outcome, and can be used as a prognosis marker for AML comparable with the known LSC score, *LSC17* (39). For this, we adopted a least absolute shrinkage and selection operator (LASSO) linear regression analysis (39–41) on genes within KDM4A regulated GEPs (*KDM4A* KD Log_2_FC ≥1 or ≤-1; adj. *p*≤0.05; supplemental file) discerning which genes best related to patient overall survival in a training subset first, and then performed regularisation in order to make the model robust to overfitting. Using this approach, a KDM4A-associated gene expression signature (GES) was constructed and calculated as the sum of the weighted expression of each of the identified 9 genes, termed *KDM4A-9* (Fig. 6D). Strikingly, high *KDM4A-9* scores were highly associated with poor overall survival (OS) in a number of large independent AML cohorts (Figs. 6E-6H) independent of age, cytogenetic risk score and frequent mutation status of known prognostic value. The robust prognostic value of the *KDM4A-9* score across diverse AML genotypes indicates, that the score may be related to the important biological activities of AML-LSCs. We find that the *KDM4A-*9 score correlates with the LSC-based biomarker*, LSC17* score of AML samples and over 75% of *KDM4A-9* high score (above medium value) fractions are LSC+ (Fig. 7A); akin to the *LSC17, KDM4A-9* is a strong predictive indicator of AML LSC activity (Fig. 7B). Interestingly, there is no overlap between these two gene signatures, prompting us to test whether a combined signature termed *KDM4A-9/LSC17*, which is calculated as the linear sum of the two LSC-related scaled (Min-max scaling) scores could further improve prediction of stemness in AML samples by ROC curve analysis. Individually, we find that the *KDM4A-9* is capable of higher sensitivity (*KDM4A-9*: 85.5% *vs. LSC17*: 68.1%) whilst the *LSC17* has better specificity (*LSC17*: 73% *vs*. *KDM4A-9*: 51.7%). Together, the combined score achieves an optimal balance between specificity and sensitivity (specificity: 67.4%, sensitivity: 76.8%) (Fig. 7C) overcoming the limitations of either the *LSC17* or the *KDM4A-9* alone. These data demonstrate that the *KDM4A-9/LSC17* combined signature score are superior LSC biomarkers, reliably predicting LSC activity in AML cells. Indeed, we observe an improvement of the combined score over the *LSC17* score’s ability to predict survival in the Beat AML dataset (Figs. 7D & 7E).

**Figure 7.**
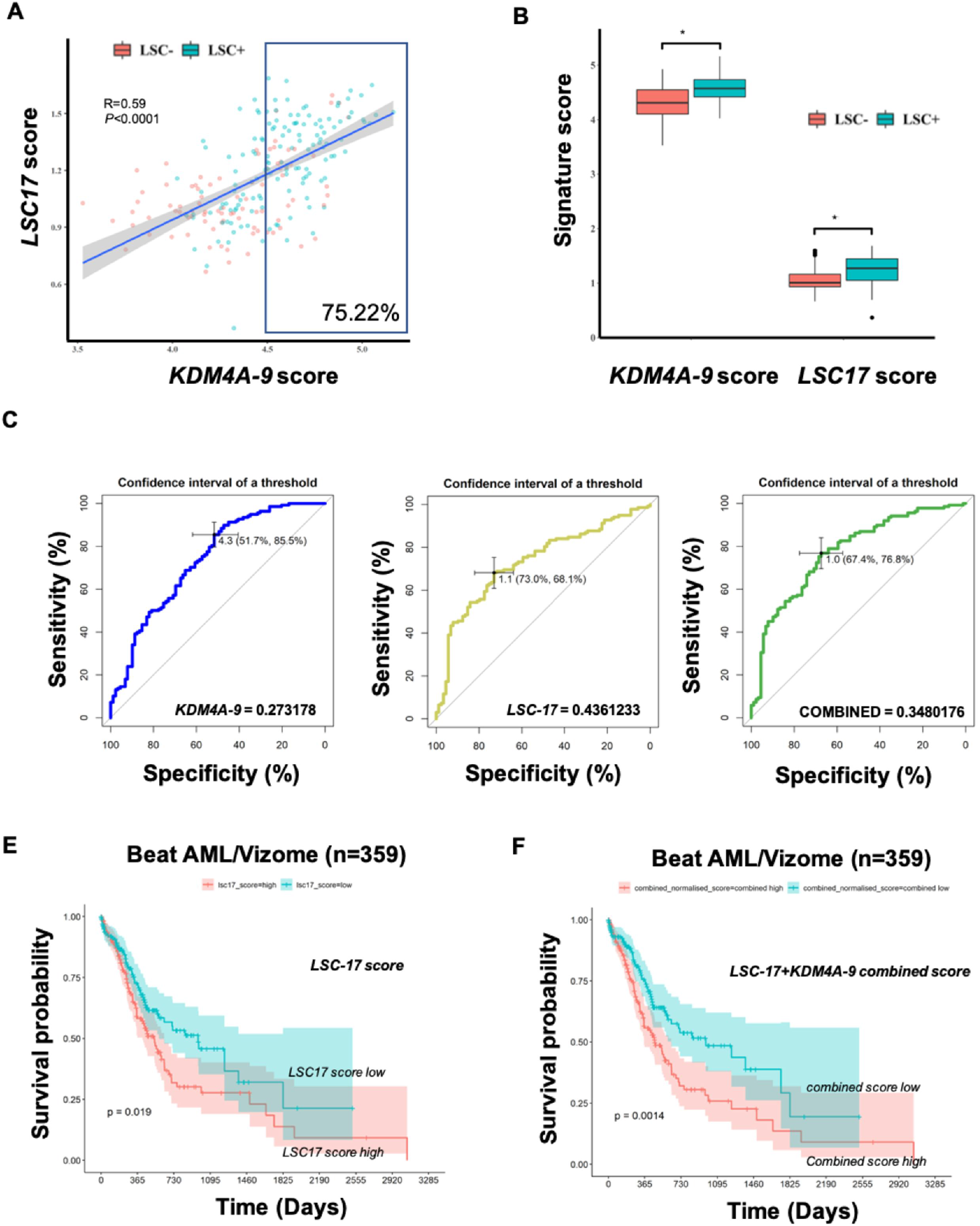
*KDM4A-9* enriched with LSC activity, is a poor prognosis marker for AML. Scatterplot showing moderate correlation between the *KDM4A-9* score and *LSC17* score in primary AML patient samples (GSE76008). LSC enriched (LSC+, n=138) cell fractions from 78 patient samples are coloured blue whilst those that lack LSC enrichment (LSC-, n=89) are coloured red. Over 75% of *KDM4A-9* high score (above median value) fractions are LSC+. Pearson correlation used to assess correlation. Significance determined by t-test. (B) Box plot showing *KDM4A-9* or *LSC17* signature scores in two comparative groups: LSC+ and LSC-from (A); unpaired *t*-test, \**p*<0.0001. (C) ROC curves of *KDM4A-9* (blue), *LSC17* (yellow) and *KDM4A-9/LSC17* (green) show the diagnostic capability of each signature to predict LSC enrichment in AML samples. The black bars in each plot are the 95% confidence intervals for the optimal cut-off. The Youden index was used to determine the optimal cut-off for each signature. (D-E) Patients in the Beat AML/Vizome dataset were dichotomized into high and low groups based on whether they possessed a score above or below the median signature score. Kaplan-Meier survival analysis conducted showing that the combined *KDM4A-9*/*LSC17* score (E) is effective in prediction of AML patient survival over *LSC17* score alone (D).

KDM4A binds at the promoter regions of *KDM4A*-9 genes, where there is enrichment of H3K9me3 and H3K27me3 signal upon *KDM4A* KD (Fig. 8A; supplemental file), suggesting a direct regulation of their expressions. We further validated these genes as *KDM4A-PAF1* axis downstream transcriptional targets by QPCR in a number of human AML cell lines and primary AML blasts following *KDM4A* KD or *PAF1* KD (Fig. 8B). Additionally, in patient AML cohorts we observed that the majority of *KDM4A-9* genes show statistically significant correlation with *KDM4A & PAF1* expressions (Figs. 8C & 8D). Furthermore, we performed weighted gene correlation network analysis (WCGNA) (42) using gene expression data from 262 diagnostic AML samples from the Beat AML dataset to evaluate the pairwise relationship between *KDM4A*, *PAF1* and the *KDM4A-9* and *LSC17* gene signatures across AML. Here, we observed that these genes have high topological overlap (topological overlap matrix (TOM) ≥0.05)) and that KDM4A represents a highly connected node within this gene network. These data demonstrate that KDM4A-PAF1 regulates the *KDM4A-9*/*LSC17* network which is persistent across AML subtypes (Fig. 8E).

**Figure 8.**
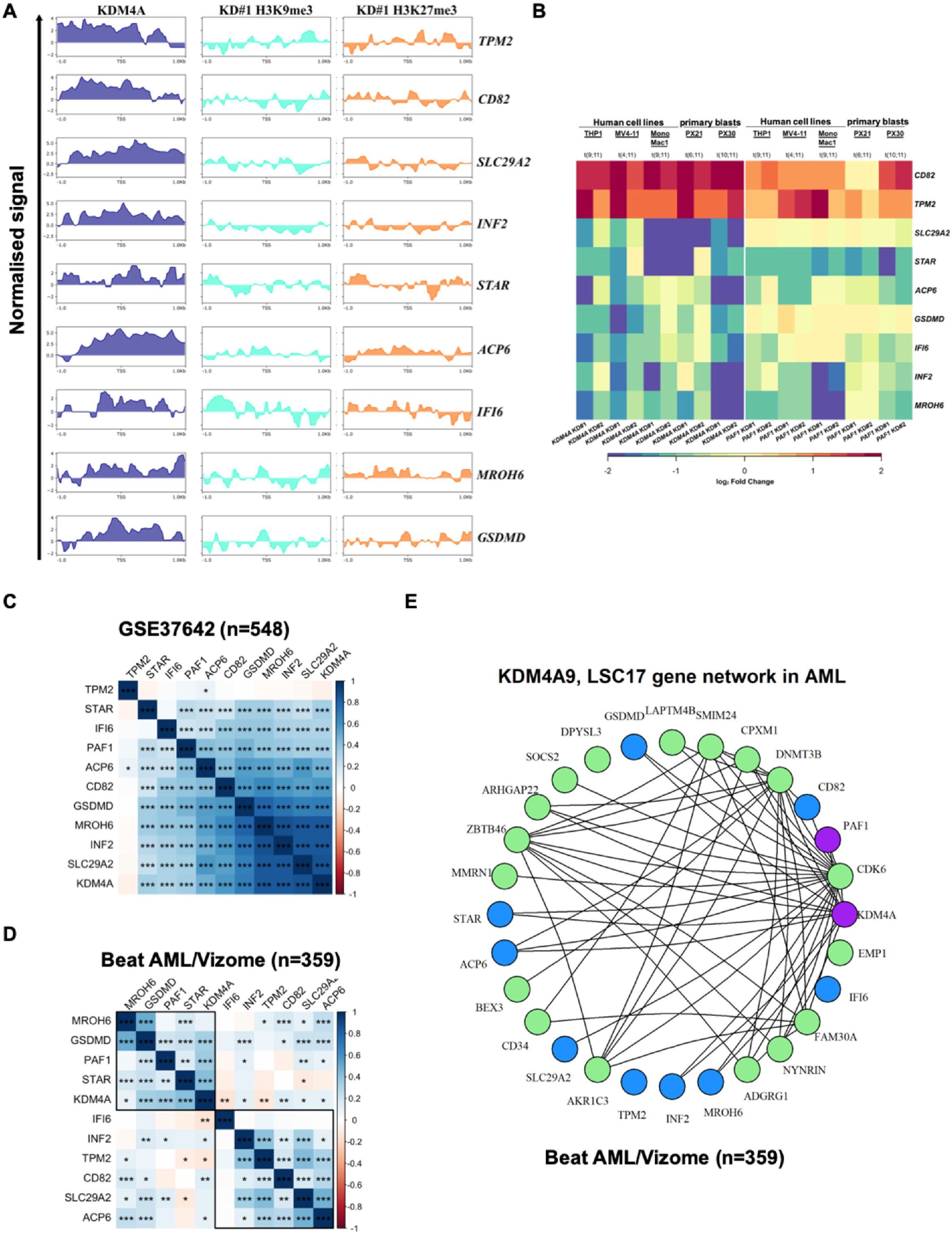
KDM4A-mediated epigenomic network required for AML cell self-renewal and survival. (A) Heatmap showing relative expression of *KDM4A-9* signature genes as determined by QPCR in the indicated human MLLr-AML cell lines and AML primary cells following *KDM4A* KD or *PAF1* KD in comparison with NTC control cells (n=3). (B) Input normalised ChIP-seq coverage tracks showing KDM4A ChIP signal in WT THP1 cells and H3K9me3/H3K27me3 ChIP signal normalized to NTC in *KDM4A* KD THP1 cells at *KDM4A-9* signature genomic loci (+/-1kb TSS). Normalised signal shown is the log_2_ ratio of read counts compared against input control. (C-D) Correlation matrices showing the Pearson correlation coefficients for *KDM4A*, *KDM4A*-9 genes and *PAF1* gene expression in GSE37642 (C) and Beat AML/Vizome (D) AML datasets. Significance determined by t-test; \**p*<0.05, \*\**p*<0.01, \*\*\**p*<0.001, \*\*\*\**p*<0.0001. (E) *KDM4A*, *PAF1*, *KDM4A-9* (in blue) and *LSC17* (in green) gene network showing the topological overlap between genes as detected from 262 AML samples (Beat AML) (the corresponding topological overlap matrix (TOM) ≥0.05 between nodes).

### KDM4A has a distinct function to another KDM4 family member, KDM4C in AML

Previously, Cheung *et al*. showed that another KDM4 family member, KDM4C is required for MLLr-AML cell survival (7), indicating an overlapping role of KDM4A and KDM4C in AML. However, forced-expression of wild-type human KDM4C failed to rescue the clonogenic activity of murine MLL-AF9 AML cells transduced with *kdm4a* KD virus (Figs. S6A & S6B), suggesting there is a role for KDM4A, that is distinct from that of KDM4C. This is also in line with the previously reported data showing no increase of global H3K27me3 level upon pharmacological inhibition of KDM4C in MLLr-AML cells (7).

Consistently, at the molecular level, *KDM4A* KD led to global transcriptional changes distinct from that of *KDM4C* KD via GSEA comparison (Fig. S6C), further supporting a unique role for KDM4A compared to KDM4C in human AML. In particular, *KDM4A* KD has no significant impact on gene expression of two established targets of KDM4C, *HOXA9* and *MEIS1* in human MLLr-AML cells. These results are also validated by Q-PCR using shRNAs targeting *HOXA9* as control (Fig. S6D). More importantly, *kdm4c* KD had no impact on *PAF1* expression, nor its associated GEP targeted by KDM4A including *KDM4A-9* and *LSC17* gene signatures (Fig. S6E). Together, these data demonstrate a unique and critical role of KDM4A in AML. supporting a KDM4A-mediated epigenomic network required for AML cell self-renewal and survival.

## Discussion

MLLr leukemia is responsible for nearly 10% of all acute leukemia with adverse prognosis; patients often develop resistance to standard chemotherapy and relapse (43). A general MLLr mechanism in AML leukemogenesis involves MLL fusion proteins associating with a number of complexes including PAF1c and DOT1Lc, leading to aberrant transcription of MLLr target genes (43, 44). Therapeutically target DOT1L has shown promising activity in preclinical studies. However, lack of tractable enzymatic activities limits the potential of PAF1 or other subunits of the PAFc as therapeutic targets. In this study, we identify a novel signalling axis of *KDM4A-PAF1* co-regulating essential oncogenic transcriptional networks is prevalent in human AML. Thus, inhibition of the histone demethylase activity of KDM4A may provide a novel alternative mean to effectively eliminate leukemic cells with broader therapeutic applications in human AML.

Previous reports indicate that the KDM4 family as a whole are required for normal hematopoiesis (8, 22) and during embryonic development (45). However, loss of individual members is well tolerated in normal cells (22). For example, *Kdm4a* is not required for embryonic stem cell function (45) and loss of a single Kdm4 family member does not grossly affect hematopoiesis (22). These data highlight the importance of identifying KDM4 family members that are selectively required for the survival of AML cells since broad inhibition of KDM4 family is associated with toxicity. Our data demonstrate KDM4A has a distinct function to another KDM4 family member, KDM4C in AML; it is selectively required for AML cell survival, with no immediate negative effect on normal hematopoiesis *in vitro*, suggesting leukemic cells are more sensitive to KDM4A depletion therefore offering a potential therapeutic window. Given the evidence that other members of the human KDM4 family are required for normal tissue development (46), our study provides a strong rationale for further development of KDM4A specific inhibitors, which are expected to have anti-leukemic properties below clinically achievable doses therefore with minimal cytotoxicity towards heathy tissues, presenting a promising strategy for novel epigenetic-based therapy in AML.

Our data support a model that KDM4A and the PAF1c cooperate to enforce an oncogenic transcriptional programme in AML cells. This is supported by recent data (6, 47) which demonstrate that PAF1c mediated recruitment of the H3K9me3 methyltransferase SETDB1 antagonizes MLL/PAF1c signalling in MLLr-AML cells and inhibition of SETB1 promotes AML cell proliferation. Consistently, our data suggest that the precise global distribution of H3K9me3 play a critical role in AML. H3K9me3 has emerged as a key player in repressing lineage-inappropriate genes, thereby impeding the reprogramming of cell identity during development and cell fate determination (27, 48). It would be interesting to determine further the clinical diagnostic/prognostic relevance of H3K9me3 in relation to KDM4A and its downstream GES in AML patients.

*KDM4A-9* shows strong therapeutic implications comparable as *LSC17* (39). A high *KDM4A-9* score distils the downstream consequences of high levels of *KDM4A-PAF1* expression in AML, probably reflecting the important biological property of KDM4A in myeloid leukemogenesis. The detailed functional relevance of the majority of *KDM4A-9* genes in leukemogenesis is largely unknown; except Tetraspanin (CD82) (49, 50) which has been suggested to play an important regulatory role in AML. Thus, further validation is needed to determine the individual function of *KDM4A-9* genes in myeloid leukemia. The association between KDM4A or its targeted transcriptional networks, such as *KDM4A-9* and known high-risk clinical features should be explored further. Given the chemotherapy-resistant phenotype of high *KDM4A-*9 score patients, these patients may better benefit more from novel molecularly targeted therapies (e.g. KDM4A-based therapy) whilst sparing low risk patients from the additional treatment related toxicity associated with intensified regimens.

## Acknowledgments

We thank Jennifer Cassels and Karen Dunn in Paul O’Gorman Leukaemia Research Centre, Glasgow, Gary Spencer, Jeff Barry, Abi Johnson and staff from the Biological Resources Unit in CRUK Manchester Institute, for technical assistance. We thank Dr. Tim Somervaille for feedbacks and critical comments on the manuscript. We thank Dr. Peter J. M. Valk (Department of Hematology, Erasmus University Medical Centre, Rotterdam) for kindly providing the survival data for the GSE6891 data set and Dr Tobias Herold and the AMLCG group for granting access to the clinical data for GSE37642.

## Authorship and conflict of interest statement

XH, MM designed the study. MEM, LM, SP, NM, RPB and AH performed the experiments. MM and XH analysed genomic data and performed the statistical analysis. XH and MM wrote the manuscript. SM, RMJL, HGJ, DV, AMM and XH provided critical support and supervised the study. All authors read and approved the manuscript. The authors have declared that no conflict of interest exists.

